# Non-linear phylogenetic regression using regularized kernels

**DOI:** 10.1101/2023.10.04.560983

**Authors:** Ulises Rosas-Puchuri, Aintzane Santaquiteria, Sina Khanmohammadi, Claudia Solís-Lemus, Ricardo Betancur-R

## Abstract

1. Phylogenetic regression is a type of Generalized Least Squares (GLS) method that incorporates a covariance matrix based on the evolutionary relationships between species (i.e., phylogenetic relationships). While this method has found widespread use in hypothesis testing via comparative phylogenetic methods, such as phylogenetic ANOVA, its ability to account for non-linear relationships has received little attention.
2. To address this issue, we utilized GLS in a high-dimensional feature space, employing linear combinations of transformed data to account for non-linearity, a common approach in kernel regression. We analyzed two biological datasets using both Radial Basis Function (RBF) and linear kernel transformations. The first dataset contained morphometric data, while the second dataset comprised discrete trait data and diversification rates as labels. Hyperparameter tuning of the model was achieved through cross-validation rounds in the training set.
3. In the tested biological datasets, regularized kernels reduced the error rate (as measured by RMSE) by around 20% compared to linear-based regression when data did not exhibit linear relationships. In simulated datasets, the error rate decreased almost exponentially with the level of non-linearity.
4. These results show that introducing kernels into phylogenetic regression analysis presents a novel and promising tool for complementing phylogenetic comparative methods. We have integrated this method into Python package named phyloKRR, which is freely available at: https://github.com/Ulises-Rosas/phylokrr.

## 1 Introduction

Phylogenetic regression is mathematically equivalent to the Generalized Least Squares (GLS), which takes into account the non-independence of data points by incorporating information from a phylogenetic tree. This is necessary because species traits do not evolve independently, as they share a common evolutionary history (Felsenstein, 1985; Harvey *et al*., 1991). For instance, the length-weight relationships can vary across different clades due to specific growth rates associated with each clade (Harvey *et al*., 1991). To incorporate phylogenetic information into the GLS, we typically infer a covariance matrix derived from species traits (Harmon, 2019). The inference of the covariance matrix assumes that the traits evolve according to a specific stochastic process, such as Brownian motion (Lefebvre, 2007), with the timing determined by the branch lengths of the phylogenetic tree. Once the covariance matrix is obtained, the data is weighted by the inverse of this matrix (see Eq. 1), and then an standard least-squares algorithm can draw a linear fit over the transformed data by finding an optimal coefficient vector *β*. This approach was initially introduced by Grafen 1989, also refered to as Phylogenetic Generalized Least-Squares (PGLS hereafter). The error function of PGLS can be expressed as:

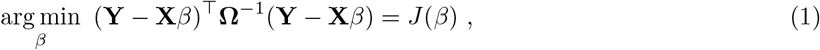

where **Y** ∈ ℝ^*n×*1^ is the response variable, **X** ∈ ℝ^*n×p*^ is the matrix with *n* taxa, or instances, with *p* number of traits, and *β* ∈ ℝ^*p×*1^ is the coefficients vector, **Ω** ∈ ℝ^*n×n*^ is the modeled covariance matrix. More justification and background for the above equation can be seen at Supporting Information S.I.

The above approach assumes that a linear fit provides the most optimal relationship between the traits and response variable. However, little attention has been given to the ability of phylogenetic regression to account for the non-linear interaction of variables in this relationship (see Fig. 1) (Quader *et al*., 2004). For instance, the length-weight relationship, which is supported by multiple lines of evidence, is known to follow to a cubic association. This relationship can be interpreted as the length variable interacting with itself in a non-linear manner. While it is possible to apply suitable transformations, like log-transformations in the context of length-weight relationships, to fit a linear function effectively, we often do not have prior knowledge about the variable interactions that drive the relationship (Harvey *et al*., 1991; Quader *et al*., 2004). Despite this limitation, the linear model remains the most widely used method in comparative biology and finds extensive application in hypothesis testing, such as phylogenetic ANOVA (Adams & Collyer, 2018).

**Figure 1:**
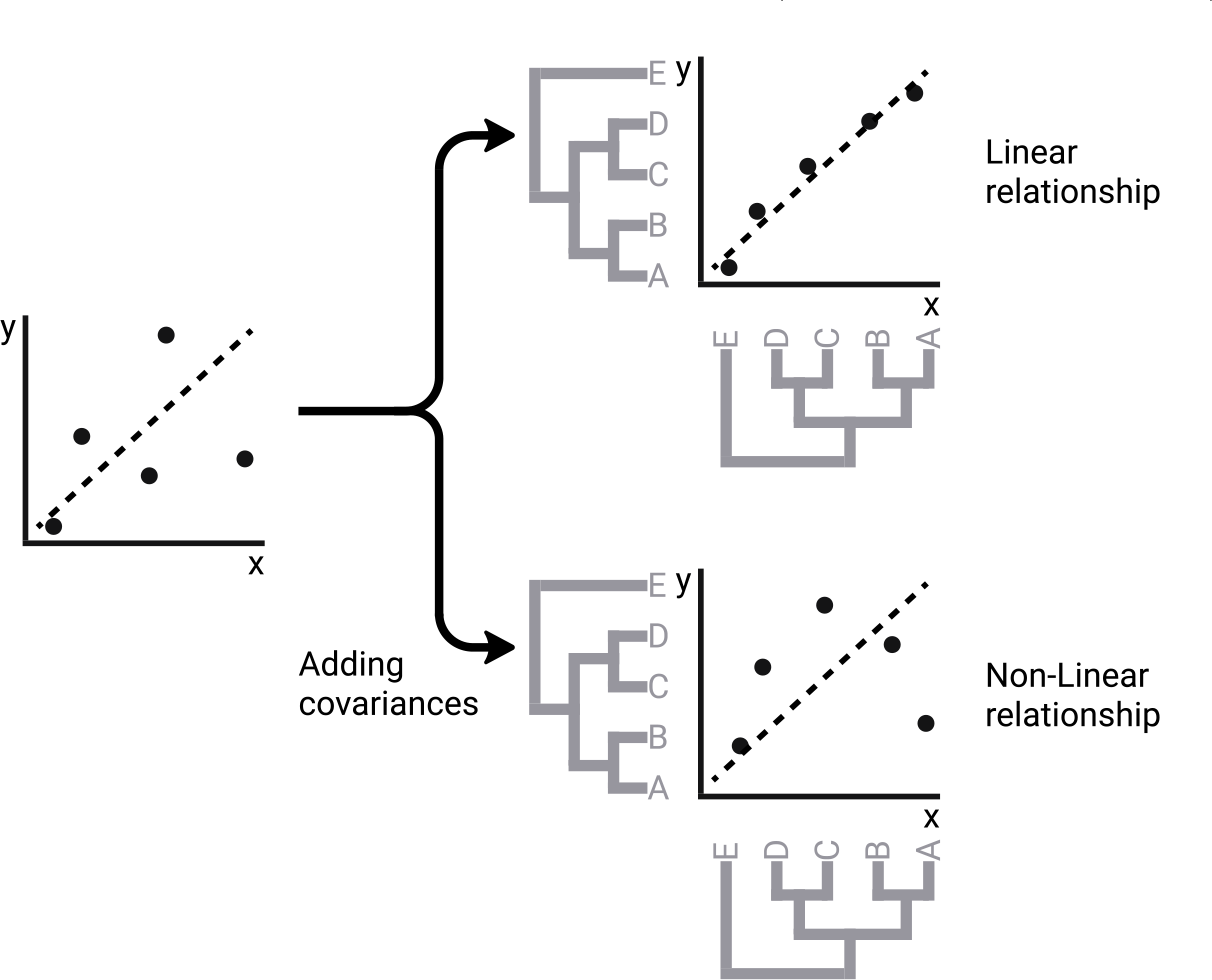
This schematic representation illustrates two possible scenarios that can occur when weighting data points with the covariance matrix to address non-independence and heteroscedasticity of the data. In the first scenario, the incorporation of covariances can effectively linearize the relationships among the data points. As a result, PGLS performs well in fitting the data, capturing the linear relationships present. Conversely, in the second scenario, even after incorporating the covariances, the data points do not linearize due to non-linear interactions between variables. In such cases, the limitations of PGLS become apparent as it may struggle to accurately capture the non-linear relationships present in the data.

While non-linear models for phylogenetic regression that use convex optimization (i.e., to minimize the error, this type of optimization assumes that every local minimum is the global minimum) have already been proposed through the use of generalized estimating equations (GEE; Martins & Hansen, 1997; Paradis & Claude, 2002; Ives & Garland Jr, 2010), they have a finite set of basis functions that are parameterized from the available exponential distribution family, which constrains the hypothesis space. On the other hand, methods like Neural Network-based models (LeCun *et al*., 2015) or decision tree-based models (Breiman, 2001; Chen *et al*., 2015) offer adaptive non-linear basis functions. However, as they are non-convex, they require substantial amounts of data to build robust models that can handle multiple local minima. An intermediate approach that combines convexity and non-linearity, and has the potential to account for an infinite number of basis functions, is kernel regression.

Kernel regression expands the number of dimensions based on linear combinations (inner products) of the original data. This expansion allows us to successfully fit a linear model in cases where it was previously unattainable. Upon projecting the resulting linear model back from this higher dimensional space to the original space, we obtain a non-linear model (Cortes & Vapnik, 1995). This technique, known as the “Kernel trick,” can be efficiently integrated into standard regression models, even when the number of additional dimensions approaches infinity (Bishop & Nasrabadi, 2006). To address the risk of overfitting, a regularization term can be added to the regression model that already employs the Kernel trick. When this regularization term is in the form of the *l*_2_-norm (see Materials and Methods), the regularized kernel is referred to as Kernel Ridge Regression (KRR) (Cristianini & Shawe-Taylor, 2000; Cortes & Vapnik, 1995). Although Regularized Kernels have found widespread use in supervised machine learning for classification models, particularly through Support Vector Machines (Cortes & Vapnik, 1995; Bishop & Nasrabadi, 2006), their potential application in phylogenetic regression remains largely unexplored. The utilization of Regularized Kernel regression is well-suited for comparative analyses due to its ability to handle a relatively small number of data features for training the model. This is particularly beneficial for biological datasets, where obtaining a large number of traits for numerous species can be costly. Additionally, Regularized Kernel regression involves only a limited number of parameters to fine-tune, typically consisting of at most two parameters.

In this study we (1) implemented a Regularized Kernel Regression model, more specifically a KRR model, that account for phylogenetic information; (2) tested two kernel functions that allowed us to expand the number of dimensions; (3) validated the obtained models in simulated data; and (4) tested the model in two biological datasets. Our main aim is to approach non-linearity in phylogenetic regressions.

## 2 Material and Methods

### 2.1 Implementation

In this section, our first step is to apply weights to the original data using the covariance matrix. By doing so, we ensure that the data incorporates the phylogenetic information. Next, we present the mathematical implementation of regularized kernel regression on this weighted data. This approach allows us to leverage the benefits of kernel regression while accounting for the phylogenetic structure present in the data.

Let **Q** and **Λ** be the eigenvectors and eigenvalues matrices of the covariance matrix **Ω**, respectively. Since **Ω** is a positive semidefinite matrix, **Q** is an orthogonal matrix. Let **P** = **QΛ**^*−*1*/*2^**Q**^T^ such that **P**^T^**P** = **Ω**^*−*1^. Then, we can re-write Eq. 1 as:

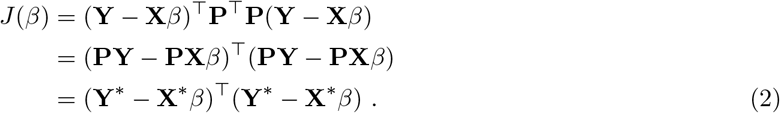

The transformed data, denoted as **Y**^*^ and **X**^*^, represents the original data weighted with the phylogenetic information derived from the **P** matrix. Moving forward, we follow the standard derivation for regularized kernel regression, as described in Bishop & Nasrabadi (2006) and Hastie *et al*. (2009).

From Eq. 2, we can add a *l*_2_-norm regularization term for *β* to control over-fitting, and increase the number of dimensions for each instance for **X**^*^ matrix as follows:

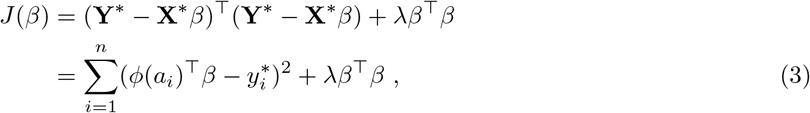

where *λ* is the regularization coefficient that controls the relative importance of the regularization term, the *a*_*i*_ is the *i*^*th*^ instance of **X**^*^, the function *φ* : ℝ^*p*^ maps the instance *a*_*i*_ into the high dimensional feature space, also known as the Reproducing Kernel Hilbert Space, and 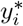 is the *i*^*th*^ element of the **Y**^*^ vector. Eq. 3 is our new error function and we can minimize it by setting set the gradient *J* (*β*) with respect to *β* equal to zero, which produces the following dual form:

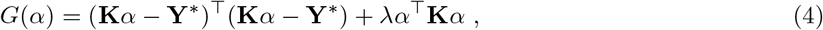

where **K** ∈ ℝ^*n×n*^ is the Gram matrix (Bishop & Nasrabadi, 2006) whose elements are inner product of instances mapped in the high dimensional feature space ℋ, and *α* ∈ R^*n×*1^ is the column vector containing elements of the form 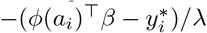. We can define each element of matrix **K** as *k*(*i, j*) = *φ*(*a*_*i*_)^T^*φ*(*a*_*j*_), where *k*(*i, j*) is the kernel function that evaluate the above dot product without explicitly mapping the instances into the high dimensional feature space ℋ. Eq. 4 is now operating in high-dimensional space and we can utilize least-squares to search for an optimal solution for *α*. This optimal solution will ultimately yield a non-linear solution to the original problem (Eq. 3), also known as the primal form. If we set the gradient of *G*(*α*) in equation 4 with respect to *α* equal to zero, we obtain:

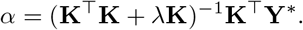

More detailed mathematical justification of the above derivations can be seen at the Supporting Information S.II. It can be shown that the time complexity of the above model is *O*(*n*^3^ + *n*^2^*p*) (Murphy, 2012), where *n* is the number of instances and *p* the number of traits. Then, even if we increase the number of dimensions possibly to infinity (see below) the time complexity is mostly influenced by the number of instances and the original number of dimensions (i.e., traits).

We tested two kernel functions, with the first one being the Radial Basis Function (RBF kernel hereafter) defined as 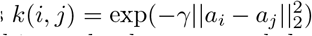. By definition, the exponential function can be expressed as a power series, and it can be demonstrated that this infinite summation results in a power series involving the inner product 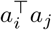 in the RBF. This property allows us to evaluate the inner product in an infinite-dimensional space. The value of *γ* controls the relative importance of the terms in the power series. For example, if *γ* is set to zero, the kernel evaluation yields a value of 1 for every inner product. When *γ* lies between 0 and 1, it is expected that the evaluation will gradually diminish in higher-order terms. However, depending on the data, even when the evaluation vanishes, it can still capture a high-dimensional space. Conversely, if the value of *γ* is too high, more weight is assigned to the higher-order terms in the power series, resulting in a possible data overfitting scenario.

The second kernel function is the linear kernel, defined as 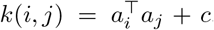. Unlike the first kernel function, this kernel directly computes the inner product without previously mapping instances into a higher-dimensional space. Although it is a simple kernel function, it proves useful in cases where the dataset already exists in a high-dimensional space, such as in the case of text classification datasets (Murphy, 2012; Géron, 2019). To introduce flexibility, we have incorporated an additional parameter *c*, which allows for adjusting the shift of the inner product.

### 2.2 Simulated datasets

For a given tree *i* with 500 species, we simulated three traits under Brownian motion (Lefebvre, 2007). The time of this diffusion process is determined by the branch lengths of the tree *i*. Each species trait is considered a realization of this process (Felsenstein, 1985), and its variance is determined by the total length of the tree *i*, while the covariance between two species is given by the sum of the branch lengths they share in the tree *i*. Let (**X**_*i*_)_:*j*_ be the column with trait *j* of the matrix containing all three traits **X**_*i*_ ∈ ℝ^500*×*3^. Then, (**X**_*i*_)_:*j*_ can have the following multivariable normal distribution:

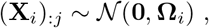

where **0** ∈ ℝ^500*×*1^ is the mean vector containing zeros, and **Ω**_*i*_ ∈ ℝ^500*×*500^ is the covariance matrix directly derived from the tree *i*. We generated 100 random coalescent trees (i.e., *i* ∈ {1, …, 100}), and extracted their covariance matrix by using the R package Ape v.5.7 (Paradis *et al*., 2004). Traits draw from the multivariable normal distribution were obtained by using the Python package Numpy v.1.21.6 (Harris *et al*., 2020).

To introduce non-linearity to the response variable, we employed the following relationship:

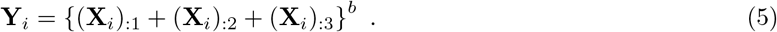

Here, *b* determines the order of non-linearity. For example, when *b* = 0, **Y**_*i*_ takes a vector of just ones. On the other hand, when *b* = 1, **Y**_*i*_ can be fully explained by a linear model. For other values of *b*, **Y**_*i*_ takes non-linear relationships, including interactions between columns of **X**_*i*_ due to the expansion of the polynomial. We tested the performance of the models (i.e., Kernel models and PGLS) over nine values of *b* ranging from -1 to 7.

### 2.3 Empirical datasets

We implemented the KRR model on two empirical datasets, each accompanied by its respective time-calibrated tree. The first dataset, referred to as the “fish dataset”, was sourced from Santaquiteria et al. (to be submitted). It comprises both discrete and continuous data from a clade of tropical and temperate marine fishes, Syngnatharia (seahorses, dragonets, goatfishes, and relatives), consisting of 323 species. Our aim was to evaluate the correlation between the geographic distributions of syngnatharian lineages and their diversification (tip) rates. We processed the categorical data from the biogeographic areas – encompassing the Indo-Pacific, eastern Pacific and Atlantic oceans – into continuous ones by using the one-hot encoding technique. This approach creates new binary columns, the count of which equals the number of categories. Each column takes a value of 1 if the species possesses the attribute related to that category, or 0 otherwise. As these binary columns represent categorical data, the data was not standardized. Though one-hot encoding is widely used in machine learning applications (Géron, 2019), this technique has also been applied in the context of phylogenetic regressions (Harvey *et al*., 1991).

The second dataset, referred to as the “primate dataset”, consists of continuous data from 90 primate species, as documented by Kirk and Kay 2004. The phylogenetic tree was obtained from the 10kTrees project (https://10ktrees.nunn-lab.org, last accessed on July 19 2023), which was reconstructed using Bayesian inference. In this case, our goal was to examine the correlation of different morphological traits with the orbit area. This data was standardized before being inputted into the model.

Assuming a Brownian motion stochastic process for all characters in both datasets, we obtained the covariance matrices using Ape v.5.7 (Paradis *et al*., 2004). The points were transformed by the covariance matrix in the response variable and trait matrix. For visualization purposes, we reduced these to one dimension using their first principal component, as both trait matrices were multivarible (Fig. 2). Notably, in the fish dataset, linear relationship between diversification rates and the PC1 of geographical distribution it is difficult to discern. Conversely, in the Primates dataset, a linear relationship is evident between Orbit area and the PC1 of morphological characters.

**Figure 2:**
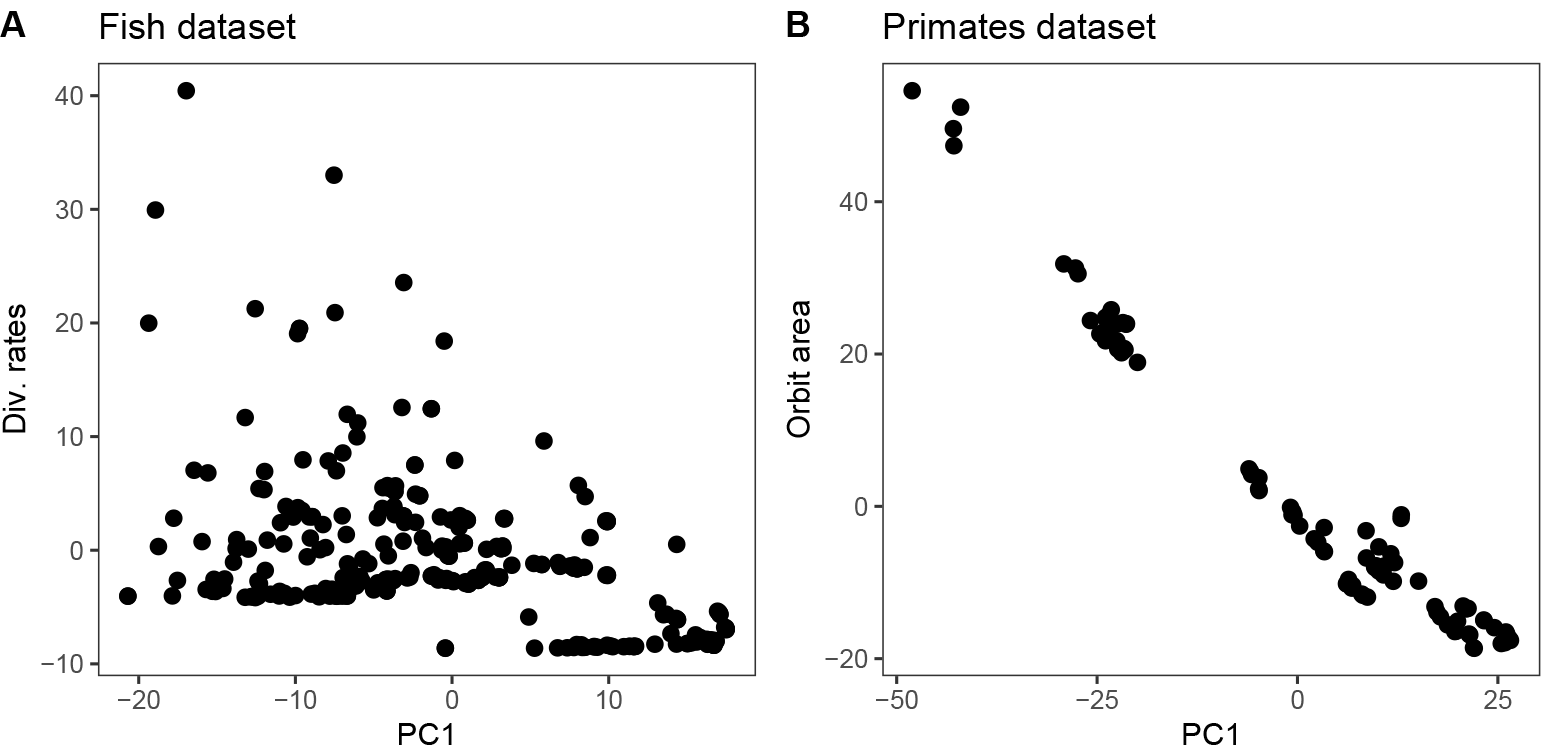
Empirical datasets. A) shows the fish dataset and B) shows primate dataset. The **X** matrix with instances and the response variable **Y** were previously transformed using phylogenetic information based on **Ω** (see Section 2.1). The x-axis in the plot above corresponds to the PC1 projection of the phylogenetic-transformed instances.

### 2.4 Evaluation

#### 2.4.1 Error estimation

To assess the performance of the model, we can divide the complete dataset into two subsets: the training set and the testing set. The training set is used to obtain the optimal *α* and corresponding hyperparameters (see below), while the testing set contains unseen data that allows for a final evaluation of the model. In the present study, the training set comprised 60% of the data, while the testing set contained the remaining 40%.

Let *α* ∈ ℝ^*n×*1^ represent the optimal solution derived from the training set, and let **Z** ∈ ℝ^*t×p*^ and **Y**_*z*_ ∈ ℝ^*t×*1^ denote the matrix of instances and the vector containing the response variable, respectively, in the testing set. Then, we can assess the model error of regularized kernel regression using the root mean squared error, RMSE_*kernel*_, as follows:

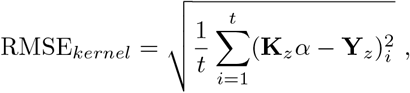

where **K**_*z*_ ∈ ℝ^*t×n*^ is the matrix where each row corresponds to the evaluated kernel function of each instance in the testing set with respect to all training instances.

#### 2.4.2 Hyperparameter tunning

We also need to determine three hyperparameters: *λ*, which adjusts the regularization process, and kernel-specific function parameters such as *γ* for the RBF kernel and *c* for the linear kernel. To accomplish this, we employed cross-validation on the training set to obtain the optimal hyperparameters. This process involves randomly splitting the training set into k-folds. Each fold serves as a test set for a particular combination of hyperparameters, while the remaining k-1 folds are used for training the model. At the end of k iterations, we obtain a set of k errors, and we report the average. We repeated the process for a set of hyperparameters and defined the optimal hyperparameter combination as the one with the lowest average error. In this study, we used 2-fold cross-validation for the primate dataset and 3-fold cross-validation for the rest of the datasets, including the simulations.

Next, using cross-validation, we randomly chose 100 hyperparameter combinations from a broad range of hyperparameters in the logarithmic scale. Specifically, we set the range of *λ* from 2^*−*5^ to 2^15^, the range of *γ* from 2^*−*15^ to 2^3^, and the range of *c* from 2^*−*5^ to 2^15^.

#### 2.4.3 Model comparison

The aforementioned approach provides us with a single optimal combination of hyperparameters. Subsequently, we can evaluate the model’s performance using the obtained *α* and hyperparameters on the testing set. To ensure robustness in the empirical datasets tests, we ran 20 rounds of cross-validation with random assignments of training and testing sets. After completing the loop, we obtained 20 sets of errors for the RBF kernel, Linear kernel, and PGLS. For each iteration where the Kernel models were tested, either in the empirical or simulated dataset, we also performed PGLS on the testing set of the same partition and assessed its error. We later compared the models with the Wilcoxon rank-sum test, where p-values were adjusted for multiple comparisons.

## 3 Results

When testing with simulated datasets, the RBF kernel model effectively captured the non-linearity of the data across various datasets, as illustrated in Fig. 3. However, when the data was linear (b = 1), both the RBF and linear kernel models showed a noticeable and statistically significant (*p <* 10^*−*34^) decrease in performance compared to PGLS. For instances where b differed from 1, the performance improvement of the RBF kernel was statistically significant (*p <* 10^*−*9^). Regarding the Linear Kernel, no significant differences were detected between it and PGLS for all tested b values (*p >* 0.05), except when b = 1 (*p <* 10^*−*33^).

**Figure 3:**
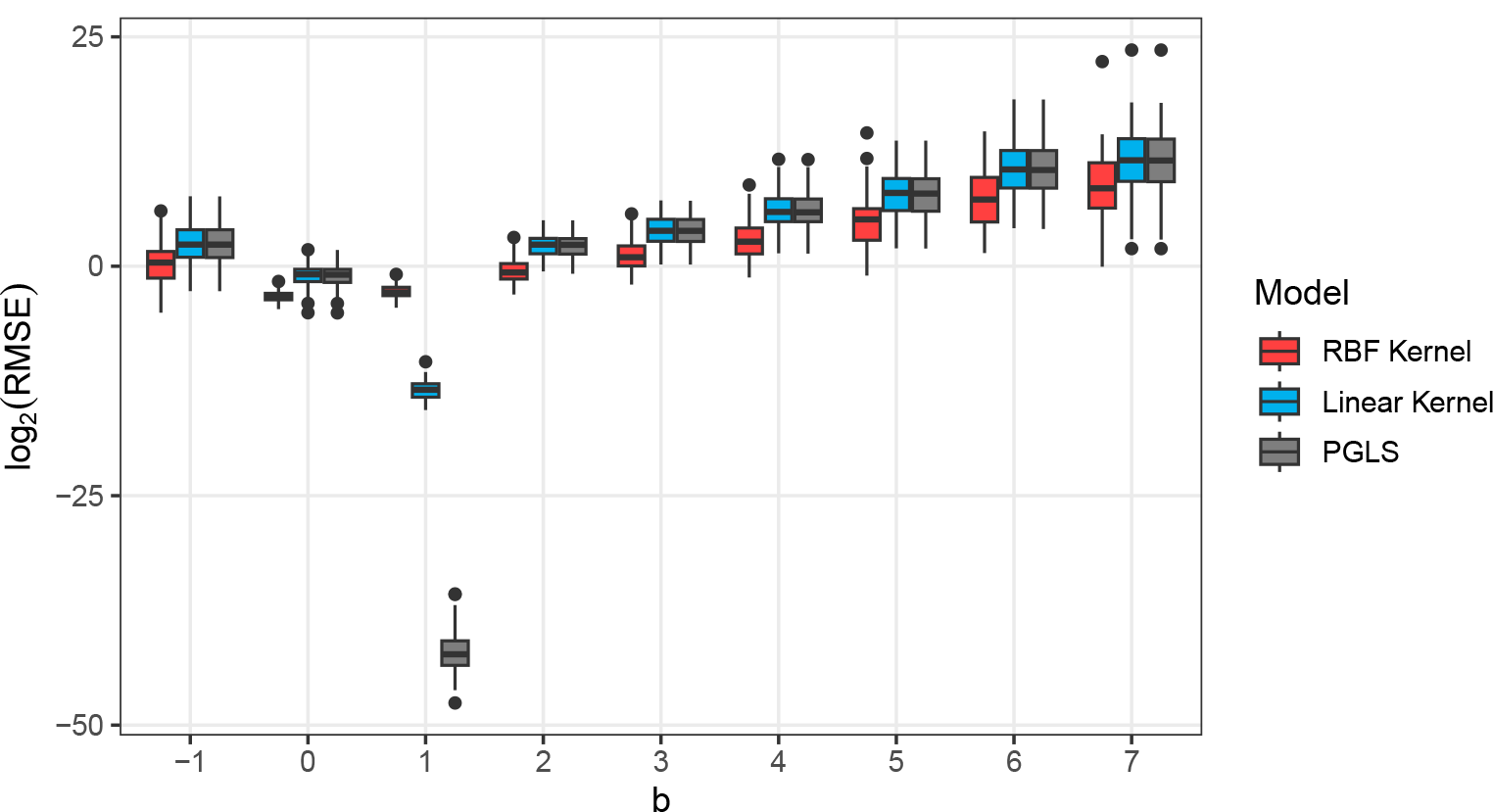
Performance of the Regularized Kernel Regression models and the PGLS under varying degrees of nonlinearity. The parameter *b* from Eq. 5 controls the nonlinearity. When *b* = 1, both Kernel models are unable to find a linear model. Each model performance at a given value of *b* used 100 simulations. We scaled the y-axis for having an improved visualization of the trend.

To assess the magnitude of the difference in the non-linear simulated datasets, we examined the difference between PGLS and Kernel model errors within each simulated dataset (Fig. 4). In the case of the RBF kernel, the minimum average difference was 0.5 for b = 0, and the maximum average difference was 10^4.94^ for b = 7. We can also observe that the difference in error between PGLS and the RBF kernel exponentially increases as the degree of non-linearity increases. For the Linear Kernel, the minimum average difference was -0.0138 for b = 0, and the maximum average difference was −10^2.61^ for b = 7.

**Figure 4:**
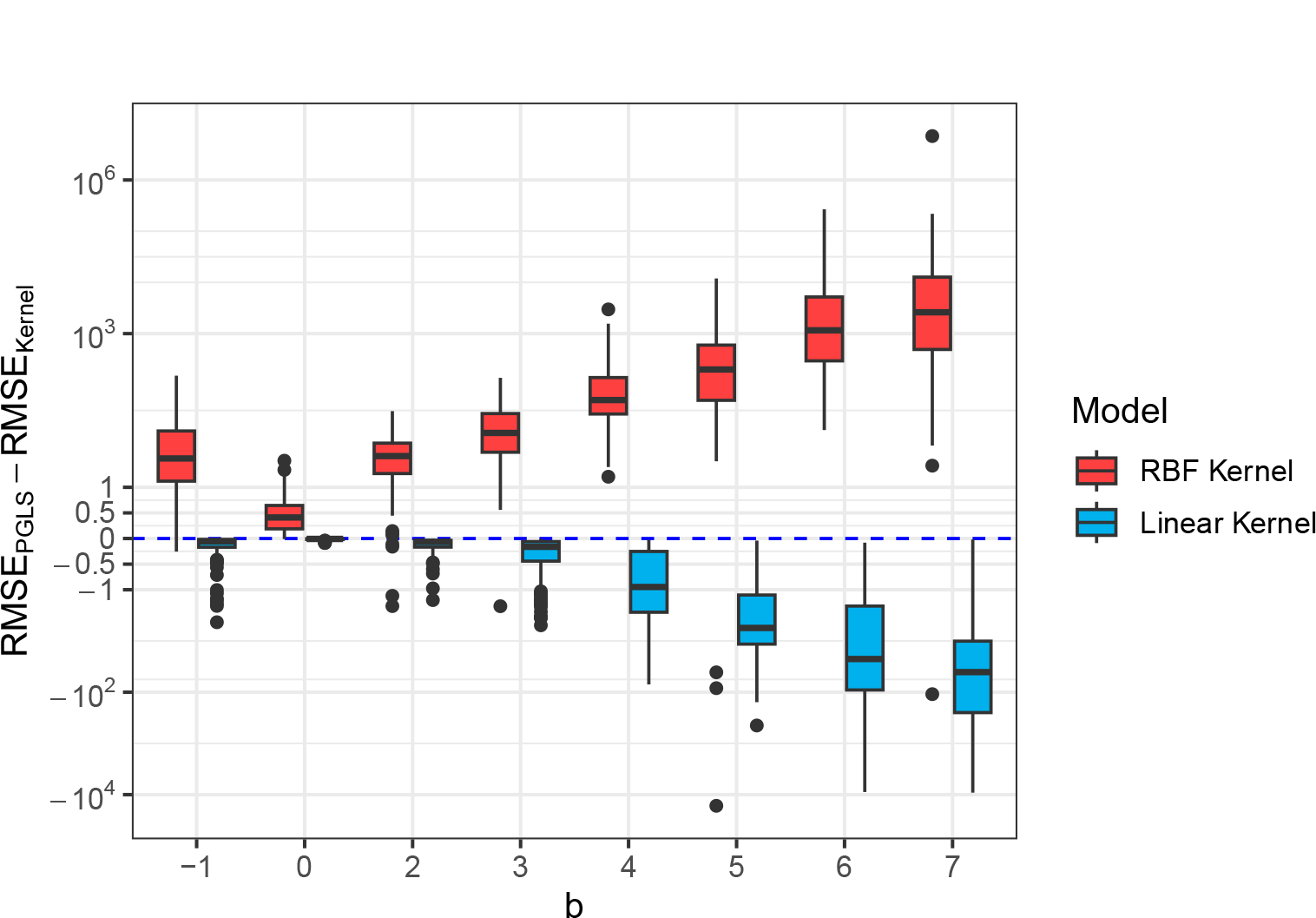
Difference between the linear model and the Regularized Kernel Regression models when the relationship is not linear. The parameter *b* from Eq. 5 controls the nonlinearity. We scaled the y-axis for having a better visualisation of the trend. The blue dashed line represents no difference between the PGLS and Regularized Kernel Regression models.

For the empirical datasets, we conducted a grid search covering all combinations of hyperparameters for a single assignment of the training and testing sets (refer to Section 2.4.3). We found that the hyperparameter space differed across the examined empirical datasets, as illustrated in Fig. 5. With the linear kernel, it became evident that the *c* hyperparameter had a minimal impact on the model performance in both datasets; the error rate remained relatively consistent beyond specific ranges of *λ* values. Based on these hyperparameter spaces, we then randomly chose the optimal hyperparameter combination for subsequent assignments of the training and testing sets.

**Figure 5:**
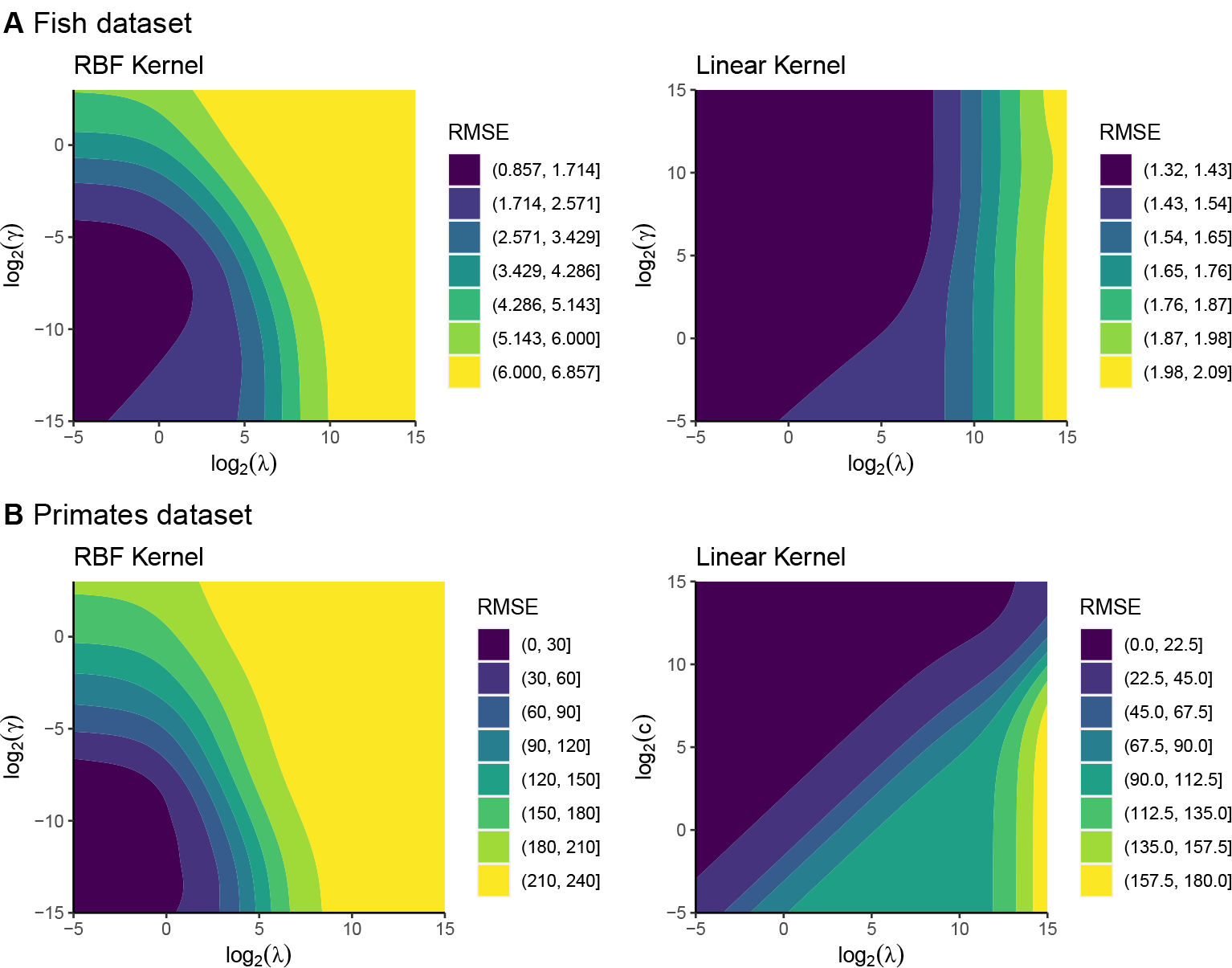
A) shows the hyperparamters space for the fish dataset using both RBF kernel and linear kernel, whereas B) shows the hyperparamter space for the primates dataset using both RBF kernel and linear kernel. The darkest area in each panel represents the space with the lowest error, as measured by RMSE. The grid search is based on one realization of the training and testing set with random assignment.

The distribution of errors is summarized in Fig. 6 for both empirical datasets. In the fish dataset, the RBF kernel exhibits a significantly superior performance compared to the Linear kernel and PGLS (*p <* 10^*−*8^ using the Wilcoxon rank-sum test). When comparing the average root mean squared error (RMSE), the RBF kernel achieves a reduction in the error of up to 20% compared to PGLS. Conversely, for the primates dataset, no statistically significant differences were observed among the tested models.

**Figure 6:**
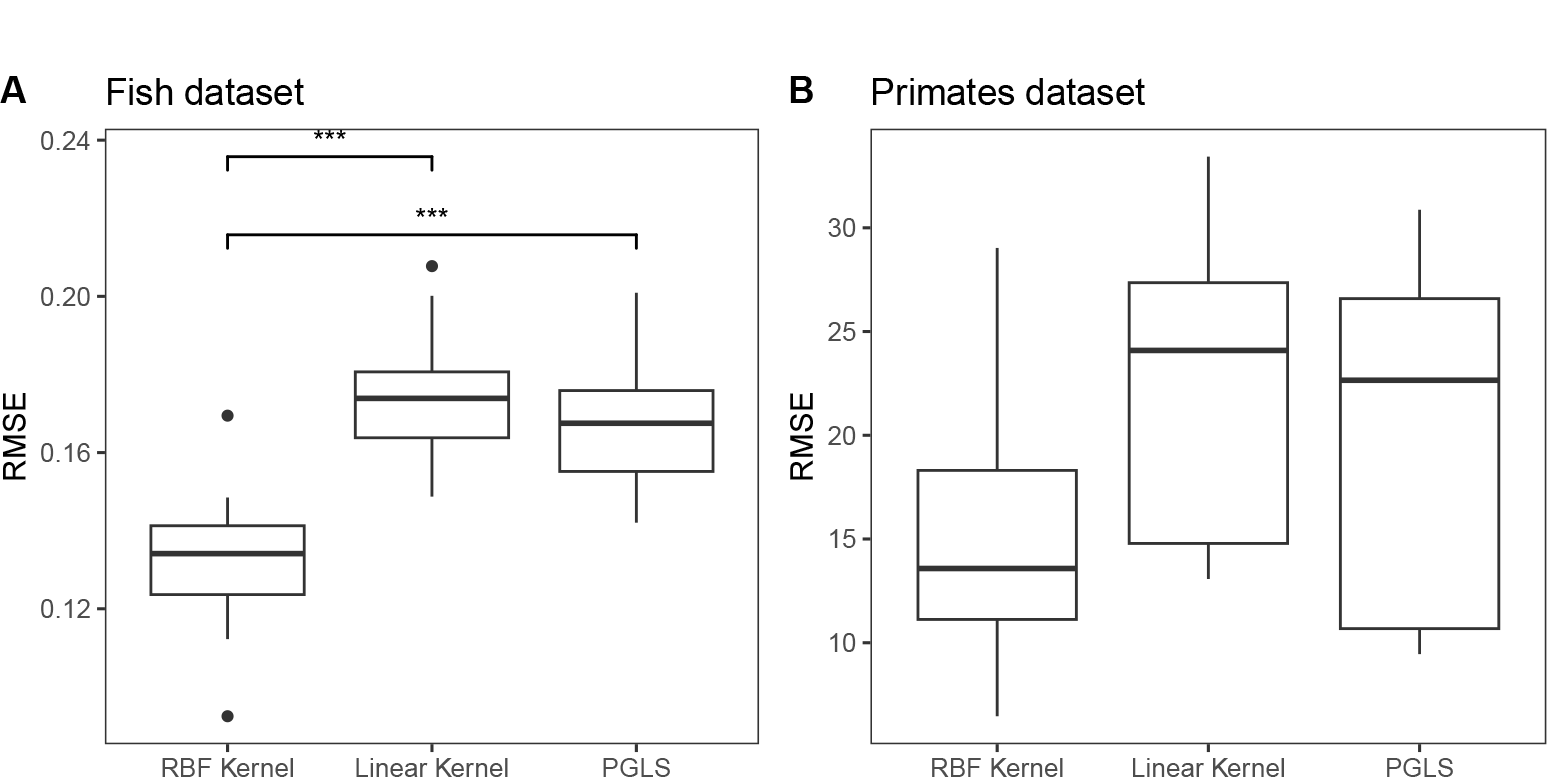
A) shows the errors of the tested models for different CV rounds. A statistically significant difference was observed between the RBF kernel and both the linear kernel and PGLS (*p <* 10^*−*8^ according to the Wilcoxon rank-sum test). B) shows the errors of the tested models for the primate dataset. In this case, no statistically significant differences were found among the models.

## 4 Discussion

In order to account for non-linearity in phylogenetic regression, we implemented regularized kernel regression incorporating phylogenetic information. Our results show that Regularized Kernel Regressions using the RBF Kernel outperform PGLS when a linear model is not the best fit. This performance advantage was apparent in both simulated and empirical datasets. However, when the datasets displayed linear behavior, the kernel regression did not surpass the PGLS, which assumes a linear model. These results show that introducing kernels into phylogenetic regression analysis presents an novel and promising tool for complementing phylogenetic comparative methods.

Machine learning models can capture the non-linearity of data efficiently, especially in the context of large datasets. They have found use in various phylogenetic analyses (e.g., Voznica *et al*., 2022; Azouri *et al*., 2021; Jiang *et al*., 2022). However, these methods rely on the assumption that samples are independent and identically distributed (Bergstra & Bengio, 2012; Vovk, 2013). This assumption may not hold true if the instances show high covariance. As a result, it is essential to use phylogenetic information to weight data, as demonstrated in Eq. 2, before feeding it into machine learning models. This step ensures the proper implementation of non-linear models that handle multi-species information data. Additionally, a similar approach as outlined in the methods section (Eq. 3), can be employed using generalized estimated equations (Eq. 3 from Paradis & Claude, 2002) to address both binary and count response variables.

The use of a regularization term in the error function enhances the model’s generalizability and robustness (Tibshirani, 1996; Zhang *et al*., 2020; Jiang *et al*., 2022), provided it is properly tuned (Srivastava *et al*., 2014; Zhang *et al*., 2021). This technique, which also assumes the independence of instances, has recently been proposed for PGLS (Adams *et al*., 2022). Even though we used a random search for hyperparameter tuning, this approach does not seem to diverge significantly from a complete grid search of hyperparameters (Bergstra & Bengio, 2012), suggesting that a fully regularized model could still be achieved. The variability in hyperparameter space across datasets underscores the importance of hyperparameter tuning when using kernels, as is also the case with other machine learning models that require such tuning (Krizhevsky *et al*., 2012).

Unregularized kernels that account for phylogenetic information have already been implemented in the context of microbiome analyses (Zhao *et al*., 2015; Xiao & Chen, 2017). However, these studies primarily focused on improving or accounting for non-linearity in the UniFrac distance (Lozupone & Knight, 2005), which is a useful metric for conducting association studies by incorporating patient traits to make predictions about diseases. Nevertheless, the use of kernels was limited to microbial abundance traits or presence/absence data, while the remainder of the model remained linear. Therefore, since the rows of the data primarily represent the same species (i.e., humans), it can be considered that it is not a true phylogenetic regression, as a phylogeny is not involved in all variables in the model.

In the case of the simulated datasets, we noticed exponentially increasing performance primarily due to the RBF kernel operating in an infinite-dimensional space. This is why we also saw high performance in the fish dataset. However, we did not observe such performance with the linear Kernel. This might be because we simulated few traits, and the linear kernel is most effective with a large number of traits, where the relationship already exists in high dimensionality (Hsu *et al*., 2003; Murphy, 2012). Although the RBF kernel converges to the linear kernel as *γ* approaches 0 (Keerthi & Lin, 2003), directly employing the linear kernel — when there is an assumption that the data is inherently high-dimensional — can be more computationally efficient. This is because there is no need for the exponentiation step to derive the complete Gram matrix.

One analytical implementation for kernels is the use of subkernels to obtain a complete kernel (Bishop & Nasrabadi, 2006). This strategy becomes particularly valuable when traits fall under different categories. For instance, in mammals, the evolution of the head bone structure may diverge from that of the rest of the body (Goswami *et al*., 2022), given the unique functions the head performs. However, determining the specific weights in a standard phylogenetic regression model for both head traits and rest-of-body traits presents challenges, especially when these traits interact in a non-linear manner. In such cases, the implementation of subkernels can be beneficial. For instance, we can define the following kernel:

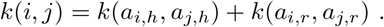

Here, *a*_*i,h*_ represents the subset of traits from instance *a*_*i*_ that belongs to the category *h* (e.g., head structures), and *a*_*i,r*_ represents the subset of traits from instance *a*_*i*_ that belongs to the category *r* (e.g., the rest of the bone structure). It is worth noting that the trait compositions for each subkernel do not need to be disjoint (Bishop & Nasrabadi, 2006). Moreover, there are several alternative methods to combine kernels, facilitating an analytical approach to understanding the structure of the data (Bishop & Nasrabadi, 2006; Duvenaud, 2014).

The incorporation of domain knowledge into the analytical model, as outlined above, provides kernel regression with an advantage over other machine learning models, such as random forests or neural networks, which also account for non-linearities. However, when handling large datasets, the process of inverting the Gram matrix in kernel regression (see Section 2.1) can become computationally demanding. In such cases, one might consider faster implementations designed for larger problems, such as the aforementioned machine learning models. Alternatively, one could select the first *m* instances to compute the Gram matrix. It has been observed that this method does not significantly compromise the accuracy of the solution (Williams & Seeger, 2000).

This study operates under the assumption that the inferred tree topology and its corresponding modeled covariance matrix are correct. However, this may not always be true if phylogenetic estimations are affected by biological processes or methodological factors (Kumar *et al*., 2012; Solís-Lemus & Ané, 2016; Kapli *et al*., 2020). Moreover, the stochastic process underlying trait evolution could potentially be misspecified (Harmon *et al*., 2010; Martin & Richards, 2019). In linear models, we have the option to use Pagel’s *λ*, which allows for adjustments to the off-diagonal values of the covariance matrix (Pagel, 1999; Revell, 2010). This provides a potential method to relax the rigidness of the covariance matrix. An alternative approach could involve the use of a set of trees (Garamszegi, 2014; Rincon-Sandoval *et al*., 2020; Santaquiteria *et al*., 2021), allowing for the testing of a range of covariance matrices. However, the challenge of incorporating covariance information into machine learning models and permitting it to evolve during the training process remains an open problem (but see McElreath, 2018, Chapter 14).

Given that kernel regression proved effective when the data was not linear, this study can serve as a baseline for future implementations of non-linear phylogenetic regressions using machine learning and complement current tools for comparative analyses. Kernels methods can effectively incorporate domain knowledge to model datasets relationships (Bishop & Nasrabadi, 2006). However, greater efforts are required to consider data non-linearity in biological analyses using machine learning models (Sapoval *et al*., 2022). We have encapsulated the model presented in this study in a Python package called phyloKRR, which is freely accessible on GitHub at: https://github.com/Ulises-Rosas/phylokrr. We have also provided a user-friendly tutorial for the package and added functions to improve the transparency of the model, including feature importance analyses (Fisher *et al*., 2019) and partial dependence plots (Elith *et al*., 2008).

## Acknowledgements

This project was funded by the National Science Foundation (NSF) under grant numbers DEB-1932759 and DEB-2225130, awarded to R.B. We appreciate a suggested reading list from Daniel Becker, which enriched our model code development and discussion section. We also thank Mehrnaz Afkhami for insightful comments during the writing process.

## Data availability

Both simulated and empirical datasets used throughout this study can be found here: https://github.com/Ulises-Rosas/phylokrr

