## Supporting Information for "Non-linear phylogenetic regression using regularized kernels"

### S.I Phylogenetic Generalized Least Squares

In this section we review results obtained from Grafen (1989); Garland and Ives (2000) for deriving the phylogenetic linear model, and give the mathematical justification for the usage of the $\mathbf{P}$ matrix for data transformation. Let the prediction of a linear model be affected by a random noise $\varepsilon \in \mathbb{R}^{n\times 1}$ that is not normally distributed:

$$\mathbf{Y} = \mathbf{X}\beta + \varepsilon \ , \tag{S1}$$

where $\mathbf{Y} \in \mathbb{R}^{n\times 1}$ is the response variable, $\mathbf{X} \in \mathbb{R}^{n\times p}$ is the matrix with instances, and $\beta \in \mathbb{R}^{p\times 1}$ is the coefficients vector.

Assume there exist an invertible matrix $\mathbf{P} \in \mathbb{R}^{n\times n}$ such that its multiplication into both sides of equation S1 generates a new random noise $\mathbf{E} \in \mathbb{R}^{n\times 1}$ that follows a multivariable normal distribution with mean vector $\mathbf{0} \in \mathbb{R}^{n\times 1}$ and covariance matrix $\sigma^2\mathbf{I} \in \mathbb{R}^{n\times n}$ :

$$\begin{aligned} \mathbf{E} &= \mathbf{P}\varepsilon = \mathbf{P}(\mathbf{Y} - \mathbf{X}\beta) \ , \\ \mathbf{E} &\sim \mathcal{N}(\mathbf{0}, \sigma^2\mathbf{I}) \ . \end{aligned} \tag{S2}$$

We can re-write equation S2 as $\mathbf{Y} = \mathbf{X}\beta - \mathbf{P}^{-1}\mathbf{E}$.

### S.I.1 Finding distribution parameters of $\mathbf{Y}$

We can estimate the new covariance matrix and mean vector of $\mathbf{X}\beta - \mathbf{P}^{-1}\mathbf{E}$ from the distribution of $\mathbf{E}$. For the $-\mathbf{P}^{-1}\mathbf{E}$ term, the mean vector and covariance matrix would be:

$$\begin{aligned}
\mathbb{E}[-\mathbf{P}^{-1}\mathbf{E}] &= -\mathbf{P}^{-1}\,\mathbb{E}[\mathbf{E}] = \mathbf{0}\ , \\
\mathrm{Cov}[-\mathbf{P}^{-1}\mathbf{E}] &= \mathbb{E}[(-\mathbf{P}^{-1}\mathbf{E} - \mathbb{E}[-\mathbf{P}^{-1}\mathbf{E}])(-\mathbf{P}^{-1}\mathbf{E} - \mathbb{E}[-\mathbf{P}^{-1}\mathbf{E}])^\top] \\
&= \mathbb{E}[(-\mathbf{P}^{-1})(\mathbf{E} - \mathbb{E}[\mathbf{E}])(\mathbf{E} - \mathbb{E}[\mathbf{E}])^\top(-\mathbf{P}^{-1})^\top] \\
&= \mathbf{P}^{-1}\mathrm{Cov}[\mathbf{E}](\mathbf{P}^{-1})^\top \\
&= \mathbf{P}^{-1}\sigma^2\mathbf{I}(\mathbf{P}^{-1})^\top \\
&= \sigma^2\mathbf{P}^{-1}(\mathbf{P}^{-1})^\top\ ,
\end{aligned}$$

without loss of generality, notice that we can define $\mathrm{Cov}[\varepsilon]$ as:

$$\begin{aligned}
\mathrm{Cov}[\varepsilon] &= \mathrm{Cov}[\mathbf{P}^{-1}\mathbf{P}\varepsilon] \\
&= \mathrm{Cov}[\mathbf{P}^{-1}\mathbf{E}] \\
&= \sigma^2\mathbf{P}^{-1}(\mathbf{P}^{-1})^\top\ .
\end{aligned} \tag{S3}$$

likewise, we can define $\mathbb{E}[\varepsilon]$ as:

$$\begin{aligned}
\mathbb{E}[\varepsilon] &= \mathbb{E}[\mathbf{P}^{-1}\mathbf{P}\varepsilon] \\
&= \mathbb{E}[\mathbf{P}^{-1}\mathbf{E}] \\
&= \mathbf{P}^{-1}\,\mathbb{E}[\mathbf{E}] = \mathbf{0}\ .
\end{aligned} \tag{S4}$$

For the complete equation $\mathbf{Y} = \mathbf{X}\beta - \mathbf{P}^{-1}\mathbf{E}$, the mean vector and covariance matrix would be:

$$\begin{aligned}
\mathbb{E}[\mathbf{X}\beta - \mathbf{P}^{-1}\mathbf{E}] &= \mathbf{X}\beta + \mathbb{E}[-\mathbf{P}^{-1}\mathbf{E}] \\
&= \mathbf{X}\beta\ , \\
\mathrm{Cov}[\mathbf{X}\beta - \mathbf{P}^{-1}\mathbf{E}] &= \mathrm{Cov}[-\mathbf{P}^{-1}\mathbf{E}] \\
&= \sigma^2\mathbf{P}^{-1}(\mathbf{P}^{-1})^\top
\end{aligned}$$

and its distribution is

$$\mathbf{Y} = \mathbf{X}\beta - \mathbf{P}^{-1}\mathbf{E} \sim \mathcal{N}(\mathbf{X}\beta, \sigma^2\mathbf{P}^{-1}(\mathbf{P}^{-1})^\top)\ .$$

Let the covariance matrix of $\mathbf{X}$ be $\mathbf{\Omega} \in \mathbb{R}^{n\times n}$, where $\mathbf{\Omega}$ is symmetric matrix and, thus, positive semi-definite matrix. Let $\mathbf{P} = \mathbf{Q}\mathbf{\Lambda}^{-1/2}\mathbf{Q}^\top$, where $\mathbf{Q}$ and $\mathbf{\Lambda}$ are the eigenvectors and eigenvalues matrices of $\mathbf{\Omega}$, respectively, such that $\mathbf{P}^{-1}(\mathbf{P}^{-1})^\top = \mathbf{\Omega}$. Then, the covariance of $\mathbf{Y}$ can be expressed in function of $\mathbf{\Omega}$:

$$\mathbf{Y} = \mathbf{X}\beta - \mathbf{P}^{-1}\mathbf{E} \sim \mathcal{N}(\mathbf{X}\beta, \sigma^2\mathbf{\Omega})\ . \tag{S5}$$

### S.I.2 Finding optimal $\beta$

Assume $\mathbf{Y}$ is independent and identically distributed (i.e., "i.i.d." assumption). Then, its density of equation S5 given by:

$$\begin{aligned}
p(\mathbf{Y} \mid \mathbf{X}, \beta) &= \frac{1}{(2\pi)^{n/2}}\frac{1}{|\sigma^2\mathbf{\Omega}|^{1/2}}\ \exp\left\{-\frac{1}{2}(\mathbf{Y} - \mathbf{X}\beta)^\top(\sigma^2\mathbf{\Omega})^{-1}(\mathbf{Y} - \mathbf{X}\beta)\right\} \\
\ln p(\mathbf{Y} \mid \mathbf{X}, \beta) &= \ln\left\{\frac{1}{(2\pi)^{n/2}}\right\} + \ln\left\{\frac{1}{|\sigma^2\mathbf{\Omega}|^{1/2}}\right\} - \frac{1}{2\sigma^2}(\mathbf{Y} - \mathbf{X}\beta)^\top\mathbf{\Omega}^{-1}(\mathbf{Y} - \mathbf{X}\beta) \\
\ln p(\mathbf{Y} \mid \mathbf{X}, \beta) &= c_1 + c_2 - c_3(\mathbf{Y} - \mathbf{X}\beta)^\top\mathbf{\Omega}^{-1}(\mathbf{Y} - \mathbf{X}\beta)
\end{aligned}$$

where $c_1$, $c_2$, and $c_3$ are constants from the model. From above equation we can clearly see that the maximization of $\ln p(\mathbf{Y} \mid \mathbf{X}, \beta)$ depends on the minimization of right most term:

$$\arg\max_{\beta} \ln p(\mathbf{Y} \mid \mathbf{X}, \beta) = \arg\min_{\beta} (\mathbf{Y} - \mathbf{X}\beta)^\top \mathbf{\Omega}^{-1} (\mathbf{Y} - \mathbf{X}\beta) \ .$$

From above relationship we can define our objective function as:

$$J(\beta) = (\mathbf{Y} - \mathbf{X}\beta)^\top \mathbf{\Omega}^{-1} (\mathbf{Y} - \mathbf{X}\beta) \tag{S6}$$

Expanding above equation, differentiating it with respect to $\beta$, and setting it to zero, we can obtain our optimal $\beta$:

$$\begin{aligned} J(\beta) &= \mathbf{Y}^\top \mathbf{\Omega}^{-1} \mathbf{Y} - 2\beta^\top \mathbf{X}^\top \mathbf{\Omega}^{-1} \mathbf{Y} + \beta^\top \mathbf{X}^\top \mathbf{\Omega}^{-1} \mathbf{X}\beta \\ \frac{\partial J(\beta)}{\partial \beta} &= -2\mathbf{X}^\top \mathbf{\Omega}^{-1} \mathbf{Y} + (\mathbf{X}^\top \mathbf{\Omega}^{-1} \mathbf{X} + (\mathbf{X}^\top \mathbf{\Omega}^{-1} \mathbf{X})^\top)\beta = 0 \\ &\Rightarrow \beta = (\mathbf{X}^\top \mathbf{\Omega}^{-1} \mathbf{X})^{-1} (\mathbf{X}^\top \mathbf{\Omega}^{-1} \mathbf{Y}) \end{aligned}$$

#### S.I.3 Finding optimal $\beta$ from transformed data

We can obtain the same optimal $\beta$ by transforming data with the $\mathbf{P}$ matrix. Since we can re-write $\mathbf{\Omega}^{-1}$ as $\mathbf{P}^\top \mathbf{P} = \mathbf{\Omega}^{-1}$, then the objective function (i.e., equation S6) can also can take this form:

$$\begin{aligned} J(\beta) &= (\mathbf{Y} - \mathbf{X}\beta)^\top \mathbf{P}^\top \mathbf{P} (\mathbf{Y} - \mathbf{X}\beta) \\ &= (\mathbf{PY} - \mathbf{PX}\beta)^\top (\mathbf{PY} - \mathbf{PX}\beta) \ . \end{aligned}$$

Let $\mathbf{Y}^* = \mathbf{PY}$ and $\mathbf{X}^* = \mathbf{PX}$, such that we can re-write above equation as:

$$J(\beta) = (\mathbf{Y}^* - \mathbf{X}^*\beta)^\top (\mathbf{Y}^* - \mathbf{X}^*\beta) \ ,$$

whose solution generates same optimal $\beta$ as the previous section. Furthermore, from equation S1 we can see the error can also get transformed by $\mathbf{P}$:

$$\mathbf{E}^* = \mathbf{Y}^* - \mathbf{X}^*\beta = \mathbf{P}\varepsilon \ , \tag{S7}$$

and, from equation S3, the covariance of $\mathbf{E}^*$ is the identity matrix times a constant:

$$\begin{aligned} \mathrm{Cov}[\mathbf{E}^*] &= \mathrm{Cov}[\mathbf{P}\varepsilon] \\ &= \mathbf{P}\mathrm{Cov}[\varepsilon]\mathbf{P}^\top \\ &= \mathbf{P}\sigma^2 \mathbf{P}^{-1} (\mathbf{P}^{-1})^\top \mathbf{P}^\top \\ &= \sigma^2 \mathbf{I} \ . \end{aligned}$$

Likewise, from equation S4, its mean vector is $\mathbf{0}$:

$$\mathbb{E}[\mathbf{E}^*] = \mathbb{E}[\mathbf{P}\varepsilon] = \mathbf{P}\,\mathbb{E}[\varepsilon] = \mathbf{0} \ .$$

### S.II Kernel Ridge Regression

#### S.II.1 Regularized least squares

Assume $\beta$ follows a multivariable normal distribution with mean vector $\mathbf{0} \in \mathbb{R}^{p \times 1}$ and covariance matrix $t^2\mathbf{I} \in \mathbb{R}^{p \times p}$ such that its density is given by:

$$p(\beta) = \frac{1}{(2\pi)^{p/2}} \frac{1}{|t^2 \mathbf{I}|^{1/2}} \exp\left\{ -\frac{1}{2t^2} \beta^\top \beta \right\} \ ,$$

Since the density of equation S7 is given by:

$$p(\mathbf{Y}^* \mid \mathbf{X}^*, \beta) = \frac{1}{(2\pi)^{n/2}} \frac{1}{|\sigma^2 \mathbf{I}|^{1/2}} \exp\left\{ -\frac{1}{2\sigma^2} (\mathbf{Y}^* - \mathbf{X}^*\beta)^\top (\mathbf{Y}^* - \mathbf{X}^*\beta) \right\} ,$$

then we can approach the posterior estimation of $\beta$ given $\mathbf{X}^*$ and $\mathbf{Y}^*$ as:

$$\begin{aligned}
p(\beta \mid \mathbf{X}^*, \mathbf{Y}^*) &\propto p(\mathbf{Y}^* \mid \beta, \mathbf{X}^*) p(\beta) \\
\ln p(\beta \mid \mathbf{X}^*, \mathbf{Y}^*) &\propto \ln p(\mathbf{Y}^* \mid \beta, \mathbf{X}^*) p(\beta) \\
&\propto \ln \left( \frac{1}{(2\pi)^{(n+p)/2}} \frac{1}{t^p \sigma^n} \exp\left\{ -\frac{1}{2\sigma^2} (\mathbf{Y}^* - \mathbf{X}^*\beta)^\top (\mathbf{Y}^* - \mathbf{X}^*\beta) - \frac{1}{2t^2} \beta^\top \beta \right\} \right) \\
&\propto \ln c_1 + \ln c_2 - \frac{1}{2\sigma^2} (\mathbf{Y}^* - \mathbf{X}^*\beta)^\top (\mathbf{Y}^* - \mathbf{X}^*\beta) - \frac{1}{2t^2} \beta^\top \beta
\end{aligned}$$

where $c_1$ and $c_2$ represent constants of the model. Thus, the Maximum A Posterior (MAP) estimation of $\beta$ is equivalent to:

$$\arg\max_\beta \ln p(\beta \mid \mathbf{X}^*, \mathbf{Y}^*) = \arg\min_\beta (\mathbf{Y}^* - \mathbf{X}^*\beta)^\top (\mathbf{Y}^* - \mathbf{X}^*\beta) + \lambda \beta^\top \beta = J(\beta) , \quad \text{(S8)}$$

where $\lambda$ is $\sigma^2/t^2$.

### S.II.2 Addition of kernels into regularized least squares

We can re-write the objective function $J(\beta)$ from equation S8 as the following:

$$\begin{aligned}
J(\beta) &= (\mathbf{Y}^* - \mathbf{X}^*\beta)^\top (\mathbf{Y}^* - \mathbf{X}^*\beta) + \lambda \beta^\top \beta \\
&= \sum_{i=1}^{n} (\phi(a_i)^\top \beta - y_i^*)^2 + \lambda \beta^\top \beta ,
\end{aligned} \quad \text{(S9)}$$

where $\phi(a_i)$ is the transformation $i^{th}$ instance of the $\mathbf{X}^*$ matrix into a high dimensional space, and $y_i^*$ is the $i^{th}$ element of the $\mathbf{Y}^*$ vector. Differentiating above equation, and setting it to zero:

$$\begin{aligned}
\frac{\partial J(\beta)}{\partial \beta} &= \sum_{i=1}^{n} 2(\phi(a_i)^\top \beta - y_i^*) \phi(a_i) + 2\lambda\beta = 0 \\
\Rightarrow \beta &= \sum_{i=1}^{n} -\frac{1}{\lambda} (\phi(a_i)^\top \beta - y_i^*) \phi(a_i) = \sum_{i=1}^{n} \alpha_i \phi(a_i) ,
\end{aligned} \quad \text{(S10)}$$

where the multiplier $\alpha_i$ represents $-(\phi(a_i)^\top \beta - y_i^*)/\lambda$. If we plug back equation S10 into equation S9, we can get the dual form of equation S9:

$$\begin{aligned}
G(\alpha) &= \sum_{i=1}^{n} \left( \phi(a_i)^\top \sum_{j=1}^{n} \alpha_j \phi(a_j) - y_i^* \right)^2 + \lambda \left( \sum_{i=1}^{n} \alpha_i \phi(a_i) \right)^\top \left( \sum_{j=1}^{n} \alpha_j \phi(a_j) \right) \\
&= \sum_{i=1}^{n} \left\{ \sum_{j=1}^{n} \alpha_j \phi(a_i)^\top \phi(a_j) - y_i^* \right\}^2 + \lambda \sum_{i=1}^{n} \left\{ \sum_{j=1}^{n} \alpha_i \alpha_j \phi(a_i)^\top \phi(a_j) \right\} .
\end{aligned} \quad \text{(S11)}$$

Let $\phi(a_i)^\top \phi(a_j) = k(i,j)$ , where $k(i,j)$ is the kernel function over $a_i$ and $a_j$. Let $i = 1$, such that

$$\begin{aligned}
\sum_{j=1}^{n} \alpha_j \phi(a_1)^\top \phi(a_j) &= \alpha_1 \phi(a_1)^\top \phi(a_1) + \cdots + \alpha_n \phi(a_1)^\top \phi(a_n) \\
&= \alpha_1 k(1,1) + \cdots + \alpha_n k(1,n) = k_{1:} \alpha ,
\end{aligned}$$

where $k_{i:}$ is the $i^{th}$ row of the Gram matrix $\mathbf{K} \in \mathbb{R}^{n \times n}$, and $\alpha \in \mathbb{R}^{n \times 1}$ is the column vector containing all alphas. Then, we can re-write the equation S11 as:

$$\begin{aligned} G(\alpha) &= \sum_{i=1}^{n} \{k_{i:}\alpha - y_i^*\}^2 + \lambda \sum_{i=1}^{n} \{\alpha_i k_{i:}\alpha\} \\ &= (\mathbf{K}\alpha - \mathbf{Y}^*)^\top (\mathbf{K}\alpha - \mathbf{Y}^*) + \lambda \alpha^\top \mathbf{K}\alpha \\ &= \alpha^\top \mathbf{K}^\top \mathbf{K}\alpha - 2\alpha^\top \mathbf{K}^\top \mathbf{Y}^* + \mathbf{Y}^{*\top} \mathbf{Y} + \lambda \alpha^\top \mathbf{K}\alpha \end{aligned}$$

Differentiating above equation with respect to $\alpha$, and set it to zero we have:

$$\begin{aligned} \frac{\partial G(\alpha)}{\partial \alpha} &= 2\mathbf{K}^\top \mathbf{K}\alpha - 2\mathbf{K}^\top \mathbf{Y}^* + 2\lambda \mathbf{K}\alpha = 0 \\ \Rightarrow \alpha &= (\mathbf{K}^\top \mathbf{K} + \lambda \mathbf{K})^{-1} \mathbf{K}^\top \mathbf{Y}^* \end{aligned}$$
